# Knowledge-Guided Learning with Curated Prior Genetic Biomarkers for Robust Model Interpretation

**DOI:** 10.64898/2026.01.27.702122

**Authors:** Beomsu Baek, Eunyoung Jang, Youngsoon Kim, Mingon Kang

**Affiliations:** Department of Computer Science, University of Nevada, Las Vegas, 89154, NV, USA; Department of Information and Statistics, Gyeongsang National University, Jinju, Republic of Korea

**Author notes:** Corresponding author. &.

**Keywords:** Knowledge-guided learning, Feature-informed learning, Survival analysis, Robust model interpretation

## Abstract

**Motivation:** Knowledge-guided learning offers effective and robust model training strategies in data-scarce settings by incorporating established domain knowledge, thereby enhancing generalization, robustness, and interpretability. By contrast, conventional deep learning approaches rely purely on data-driven learning, which can limit robust model interpretability, particularly in high-dimensional settings with limited size samples. In computational biology, knowledge-guided learning has primarily leveraged network- and structural-based knowledge, leading to biologically interpretable representations and enhanced predictive performance compared to conventional approaches. However, curated biomarkers, one of the most accessible forms of biological knowledge, remain largely unexplored within knowledge-guided paradigms.

**Results:** In this study, we propose a model-agnostic training paradigm, Biomarker-driven Explainable Prior-guided Learning (BioExPL), that can be applied to any neural networks that incorporates curated prior knowledge. BioExPL enforces neural networks to reflect curated biomarker priors in their latent representations through a novel knowledge-alignment loss. BioExPL consistently demonstrated significantly improved predictive performance and enhanced model interpretability with minimized computational overhead in simulation studies and intensive experiments on multiple cancer datasts. BioExPL not only integrates prior curated knowledge into the model but also accurately identifies unknown associated signals additionally. BioExPL is model-agnostic and domain-independent, enabling its integration into diverse neural network architectures.

**Availability and implementation:** The open-source is publicly available at: https://github.com/datax-lab/BioExPL.

## 1. Introduction

Recent advances in artificial intelligence has produced major breakthroughs in computational biology, delivering highly accurate modeling approaches for a broad range of applications, such as disease risk prediction (Zhang et al., 2025), survival analysis (Cui et al., 2025), and biomarker discovery (Arango-Argoty et al., 2025). Deep learning models, in particular, have shown remarkable effectiveness by leveraging highly flexible, data-driven approaches to learn intricate biological mechanisms. Despite the advances in AI, its use in biology is still severely constrained by the scarcity of data. Genomic datasets frequently contain tens of thousands of features measured from only a few hundred patients, which not only degrades predictive performance but also raises concerns about the robustness of models for both prediction and interpretation. Moreover, deep learning models that are trained from scratch face challenges in obtaining reproducible interpretations and frequently fail to identify known biomarkers when interpreting the model.

The knowledge-guided paradigm has emerged as a promising approach to mitigate these challenges. The paradigm incorporates domain knowledge into deep learning models to improve generalization, robustness, and interpretability (Zhou et al., 2025). For instance, physics-informed neural networks integrate physical laws into neural network training, resulting in substantial gains in robustness and reliability (Raissi et al., 2019). Moreover, a variety of knowledge-guided approaches have been proposed, including constraint-based learning (Lin et al., 2025), equation-regularized neural networks (Kuzhiyil et al., 2024), and structure-aware representation learning (Xu et al., 2025). These studies have demonstrated that incorporation of prior knowledge into deep learning models can markedly enhance both predictive performance and scientific credibility in various scientific domains.

In computational biology, prior knowledge can be broadly categorized into three types: network- and structural-based knowledge and curated biomarkers priors. First, network-based knowledge captures interactions among molecular entities, such as gene–gene and protein–protein interactions, and is commonly incorporated through graph-based modeling. The network-informed learning introduces relational dependencies to the models, facilitating the identification of interaction-driven biological effects (Zitnik et al., 2018). Second, structural-based knowledge, such as biological pathway databases and functional modules, were incorporated by constraining neural network architectures. For instance, pathway-informed models embed curated pathway structures directly into model connectivity to enable pathway-level interpretation, while improving robustness and biological plausibility in genomics applications (Hao et al., 2018; Elmarakeby et al., 2021). Lastly, curated biomarkers priors reduced the high-dimensional spaces of the input data by considering only validated evidence derived from experimental and population-level studies. However, curated biomarker approaches remain largely unexplored within knowledge-guided paradigms.

The most straightforward schemes that leverage curated biomarker priors are feature filtering and feature weighting (Pudjihartono et al., 2022). Feature filtering restricts model inputs to a predefined set of curated biomarkers, whereas feature weighting prioritizes curated biomarkers by assigning them greater weights relative to other features. These strategies have been widely adopted in computational biology to constrain high-dimensional molecular profiles to a biologically relevant subspace of characteristics (van Hilten et al., 2021; Kim and Lee, 2023; Liu et al., 2024). However, these schemes have inherent limitations. Penalizing non-curated biomarkers hinders the model in identifying potentially relevant signals that can be associated with the target outcome (Libbrecht and Noble, 2015). Relevant biomarkers often appear weak or irrelevant in isolation in complex biological systems, yet they become informative and predictive only when considered in combination with others or within particular biological contexts.

In this study, we propose a model-agnostic training paradigm, Biomarker-driven Explainable Prior-guided Learning (BioExPL), that incorporates curated prior knowledge into neural networks. (Fig. 1). BioExPL guides representation learning to reflect specified biologically validated biomarkers to the optimal model by aligning latent embeddings with knowledge-induced representations. This design drives models to embed curated biomarkers into their latent representations, while still allowing the exploration of uncharacterized potential biomarkers, resulting in enhanced predictive performance and robust model interpretation. We validated BioExPL’s performance in a controlled simulation setting and survival analysis experiments with multiple cancer datasets.

**Fig. 1.**
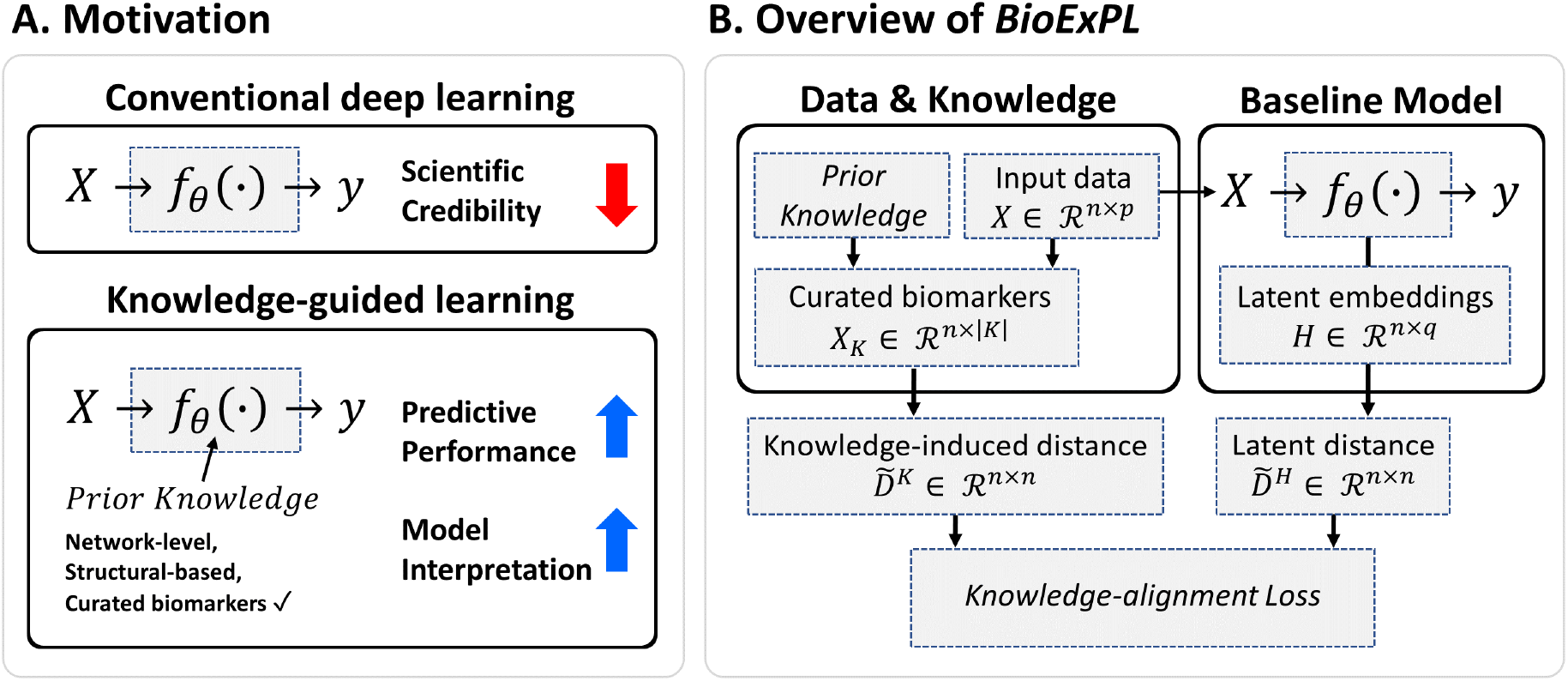
Overview of the study. (A) Motivation. Comparison between conventional deep learning and knowledge-guided learning. (B) Overview of BioExPL. A graphical representation of BioExPL, illustrating how curated biomarkers guide representation learning through a knowledge-alignment loss.

## 2. Methods

### 2.1. Objective function for knowledge-guided learning

We present BioExPL, a new knowledge-guided learning approach that serves as an effective training strategy by introducing a novel knowledge-alignment loss regularization. Let **X** ∈ ℜ^*n×p*^ denote the input data matrix, where *n* is the number of samples and *p* is the total number of features. Based on prior biological knowledge, the feature space is conceptually partitioned into two disjoint subsets: the set of knowledge-guided features **X**_*K*_, indexed by *K* ⊂ {1, …, *p*}, which are known to be associated with the target outcome, and the set of non-knowledge features indexed by the complement of *K*, denoted as **X**_*¬K*_.

Let *f*_*θ*_(·) denote a neural network baseline model with parameters *θ*, which is trained by minimizing a task-specific loss ℒ_task_ (e.g., mean squared error or cross-entropy). The objective function of BioExPL combines a task-specific loss with the knowledge-alignment loss (ℒ_KAL_):

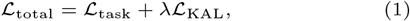

where *λ* ≥ 0 is a hyperparameter controlling the strength of the knowledge alignment. A larger value of *λ* enforces stronger alignment, thereby driving the model to emphasize knowledge-guided features **X**_*K*_. Conversely, a smaller value of *λ* allows the model to identify potential signals in non-knowledge features **X**_*¬K*_.

### 2.2. Knowledge-alignment loss

The knowledge-alignment loss is formulated by aligning two sample-wise distance matrices: (1) a knowledge-induced distance matrix 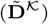 derived from **X**_*K*_, and (2) a latent distance matrix 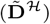 computed from intermediate latent embeddings of the neural network. The knowledge-induced distance matrix serves as a fixed reference encoding prior knowledge. The latent distance matrix is optimized to align with the knowledge-induced distance matrix during training. ℒ_KAL_ minimizes the sum of squared errors (SSE) between the distance matrices:

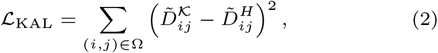

where 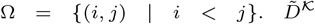 is computed solely from the knowledge-guided features **X**_*K*_. Minimizing ℒ_KAL_ therefore drives the latent embeddings to primarily embed the knowledge-guided features **X**_*K*_, while still allowing to incorporate non-knowledge features.

Given a subset matrix with knowledge-guided features, **X**_*K*_ ∈ ℜ^*n×*|*K*|^, the knowledge-induced distance matrix **D**^*K*^ ∈ ℜ^*n×n*^ is calculated by the sample-wise Euclidean distance:

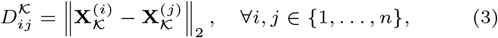

where the superscript (*i*) denotes the *i*-th sample. The latent distance matrix is derived from an intermediate mapping of the neural network. The intermediate layer can correspond to any hidden layer or an embedding layer of a neural network, *f*_*θ*_(**X**) = *g*_*ϕ*_(*h*_*ψ*_(**X**)) depending on the specific model architecture, where *h*_*ψ*_(·) represents a encoding function up to an intermediate layer and *g*_*ϕ*_(·) denotes the remaining task-specific mapping. The latent embeddings are given by

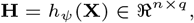

where *q* denotes the latent dimensionality. Based on the two embeddings, the latent distance matrix **D**^*H*^ ∈ ℜ^*n×n*^ is computed as

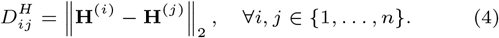

To ensure the scale comparability, both distance matrices are normalized by their respective maximum values:

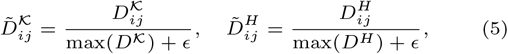

where *ϵ* is a small constant preventing division by zero. Both **D**^*K*^ and **D**^*H*^ are distance matrices with a minimum value of zero, resulting in scale-invariant matrices (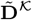 and 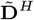) within the range [0, 1].

### 2.3. Implementation details and computational efficiency

BioExPL relies on sample-wise distance matrices, which incur a computational cost of 𝒪(*n*^2^). To mitigate this potential overhead, BioExPL is implemented in a computationally efficient manner. During mini-batch training with batch size *b*, the computation of distance alignment loss is performed only within each batch, reducing the per-iteration computational cost from 𝒪(*n*^2^) to 𝒪(*b*^2^). The knowledge-induced distance matrix **D**^*K*^ serves as a fixed reference, is computed only once prior to training, and is reused throughout optimization without further updates. BioExPL does not modify the architecture of the baseline model and is applied exclusively during training, which consequently incurs minimally additional computational for training and no memory overhead during inference.

## 3. Experimental Results

We evaluated BioExPL through simulation studies and survival analysis using multiple cancer datasets. First, the simulation study assessed whether BioExPL can effectively guide neural networks using predefined knowledge-guided features under controlled settings, where the ground truth of the true associated feature set is known. Second, we applied the proposed BioExPL to real cancer datasets to assess predictive and interpretation performance, comparing with the baseline and benchmark models.

### 3.1. Simulation Study

In the simulation study, we designed a synthetic dataset with pre-defined ground truth, where only a subset of features were defined as truly associated features (i.e., **X**_𝒯_). The simulation study informs only a subset of those truly associated features as knowledge-guided features (i.e., **X**_*K*_), instead of guiding all truly associated features, so that BioExPL can discover the un-informed set. Under this setting, we tested the following hypotheses: (1) the proposed knowledge-guided learning strategy, BioExPL, improves predictive performance compared to a baseline model; (2) BioExPL accurately identifies knowledge-guided features (**X**_*K*_) more than the baseline; (3) BioExPL effectively suppresses irrelevant features (**X**_*¬*𝒯_); and (4) BioExPL can discover truly associated features that are not included in the predefined knowledge-guided feature set (**X**_𝒯 *\K*_).

We generated an input data matrix **X** ∈ ℝ^1,000*×*100^, where each feature was independently generated from a normal distribution: **X** ∼ *N* (0, 1). Among the 100 features, we designated the first five features (i.e., **x**_1_–**x**_5_) as truly associated features, and **x**_6_–**x**_100_ as irrelevant features. The response variable was generated as a nonlinear combination of the associated features:

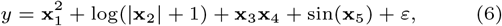

where *ε* is a noise term that follows a zero-mean Gaussian distribution and was scaled to satisfy a signal-to-noise ratio (SNR) of 10. To ensure comparable effect sizes across the associated features, each nonlinear component in (6) was scaled to have unit variance of mean zero before generating the response variable. Among the five associated features, we designated only the first three features as the knowledge-guided feature set (i.e., **X**_*K*_ = {*X*_1_, *X*_2_, *X*_3_}). Note that *x*_4_ and *x*_5_ were associated features but were not included in the knowledge-guided feature set. *x*_4_ was designed to interact with the knowledge-guided feature *x*_3_, and *x*_5_ was with an independent effect with nonlinear association. We employed a simple multilayer perceptron model (MLP) with three hidden layers as the baseline model and applied BioExPL with the hyperparameter *λ* varying over {10^−3^, 10^−2^, 10^−1^, 10^0^, 10^1^}. To implement the knowledge alignment loss, we computed the latent embeddings (**H**) from the first hidden layer of the MLP. We conducted 5-fold cross-validation and repeated the experiment 30 times, reporting the average mean squared error (MSE) across all folds and repetitions.

BioExPL consistently improved the performance of the baseline MLP (i.e., *λ* = 0) over varying *λ* (Fig. 2A). Specifically, BioExPL achieved an average MSE of 0.7374±0.0142, 0.6096±0.0115, 0.5558±0.0099, 0.5592±0.01480 and 0.6491±0.0143 at *λ* ∈ {10^−3^, 10^−2^, 10^−1^, 10^0^, 10^1^}, respectively. These results correspond to improvements of 29.7%, 41.8%, 47.0%, 46.6%, and 38.1% compared the baseline MLP (1.0490±0.0054). The improvements were further statistically validated using the Wilcoxon rank-sum test, with *p <* 0.01 for all values of *λ*.

**Fig. 2.**
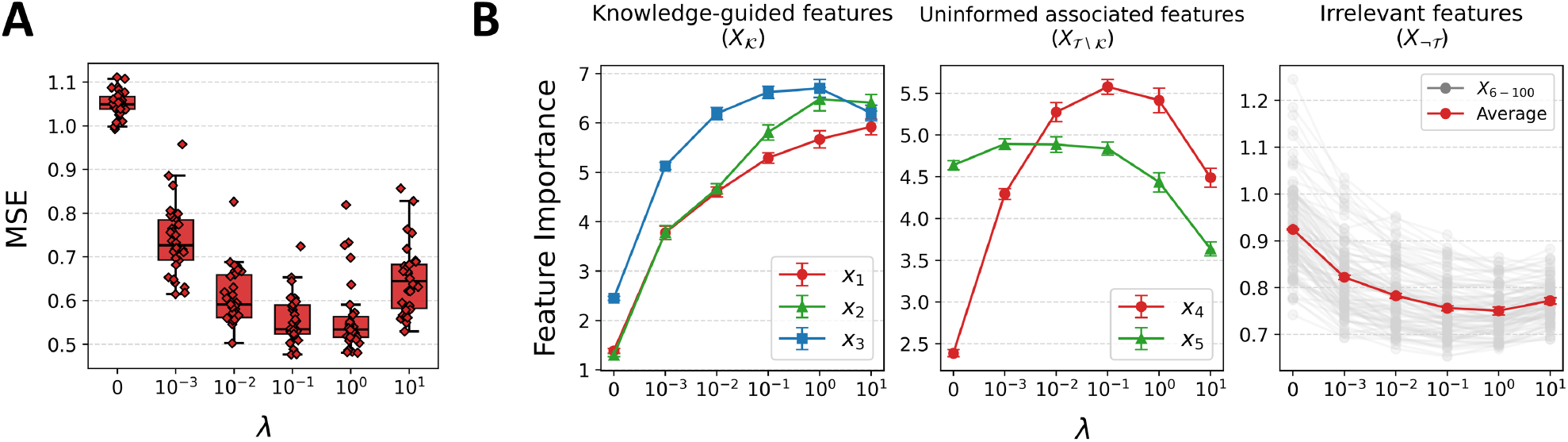
Experimental results on synthetic data. (A) Predictive performance across a range of *λ* values. Each point represents the average mean squared error (MSE) obtained from 5-fold cross-validation. (B) Feature importance across different values of *λ*. Feature importance scores were computed using a gradient-based saliency method based on input gradients, and the 1,000 feature importance values were normalized to sum to one to ensure comparability across experiments.

Furthermore, we analyzed feature importance to investigate whether BioExPL can effectively identify the predefined knowledge-guided features compared to the baseline model. We computed importance scores of each variable using a gradientbased saliency method based on input gradients for model interpretation. BioExPL with *λ* = 10^−1^, which achieved the best predictive performance, increased the average importance score of knowledge-guided features by 244.8% compared to the baseline model, from 1.7135±0.0300 to 5.9084±0.1188. In contrast, the average importance scores of irrelevant features decreased by 18.2%, from 0.9245±0.0015 to 0.7564±0.0053 (Fig. 2B, **X**_*¬*𝒯_).

We observed that BioExPL successfully identified *x*_4_ and *x*_5_ as significant features, although they were not informed to the learning (Fig. 2B). The average importance score of *x*_4_ and *x*_5_ increased by 48.2% compared to the baseline model (from 3.5136±0.0431 to 5.2075±0.0830), in contrast to the importance scores of the irrelevant features. Specifically, BioExPL was more effective at identifying features that exhibit interaction effects with knowledge-guided features. The average importance score of *x*_4_, which interacts with the knowledge-guided feature *x*_3_, increased by 133.5% (from 2.3885±0.0425 to 5.5768±0.0895), while the average importance score of *x*_5_ showed an increase of 4.3% (from 4.6387±0.0549 to 4.8384±0.0797).

### 3.2. TCGA Survival Analysis

We conducted survival analysis using cancer gene expression data to evaluate BioExPL in a real-world application. We applied BioExPL to several state-of-the-art survival analysis models across multiple cancer datasets from The Cancer Genome Atlas (TCGA). Specifically, we considered ten TCGA cancer datasets, including Liver Hepatocellular Carcinoma (LIHC), Breast Invasive Carcinoma (BRCA), Colon Adenocarcinoma (COAD), Lung Squamous Cell Carcinoma (LUSC), Skin Cutaneous Melanoma (SKCM), Lung Adenocarcinoma (LUAD), Acute Myeloid Leukemia (LAML), Kidney Renal Clear Cell Carcinoma (KIRC), Kidney Renal Papillary Cell Carcinoma (KIRP), and Glioblastoma Multiforme and Lower Grade Glioma (GBM/LGG). In this experiment, we obtained curated cancer-associated genes as the knowledge-guided feature set for each cancer type, from the comparative toxicogenomics database (Davis et al., 2025) that provides curated gene–disease associations^1^ supported by extensive prior biological literature. An brief description of the ten cancer datasets is provided in Table 1.

**Table 1.** Summary of TCGA cancer datasets.

| Cancer type | Description | Samples | Total genes | Curated genes |
| --- | --- | --- | --- | --- |
| LIHC | Liver Hepatocellular Carcinoma | 372 | 20,021 | 478 |
| BRCA | Breast Invasive Carcinoma | 1,099 | 20,194 | 466 |
| COAD | Colon Adenocarcinoma | 378 | 19,940 | 269 |
| LUSC | Lung Squamous Cell Carcinoma | 495 | 20,167 | 231 |
| SKCM | Skin Cutaneous Melanoma | 463 | 20,167 | 198 |
| LUAD | Lung Adenocarcinoma | 508 | 20,095 | 138 |
| LAML | Acute Myeloid Leukemia | 163 | 19,649 | 117 |
| KIRC | Kidney Renal Clear Cell Carcinoma | 534 | 20,194 | 113 |
| KIRP | Kidney Renal Papillary Cell Carcinoma | 290 | 20,104 | 113 |
| GBM/LGG | Glioblastoma Multiforme and Lower Grade Glioma | 693 | 19,777 | 87 |

Gene expression (RNA-seq) data were obtained from The Cancer Genome Atlas (TCGA), and curated biomarkers were defined using gene-disease associations from the Comparative Toxicogenomics Database (CTD). Note: samples denotes the number of patients, total genes denotes the number of gene features, and curated genes indicate the size of the biologically validated biomarker set used as prior knowledge. Gene expression data were obtained from TCGA, and curated gene sets were constructed based on gene-disease associations provided by the Comparative Toxicogenomics Database (CTD).

To evaluate the general applicability of BioExPL across diverse baseline architectures, we considered three survival analysis models: Cox-nnet (Ching et al., 2018) and DeepSurv (Katzman et al., 2018) have been widely used for survival analysis, and AESurv (Shen et al., 2024) was a recently proposed state-of-the-art model. Cox-nnet and DeepSurv are neural network–based proportional hazards models that estimate the log-partial hazard. Cox-nnet uses a single hidden layer, whereas DeepSurv employs a deeper architecture with multiple hidden layers. AESurv extends neural-network–based Cox models by introducing an autoencoder architecture that learns compact latent representations for survival analysis. For all the baseline models, we used the default model architectures reported in their original publications. We applied BioExPL to each model by computing the latent embeddings from the first hidden layer for Cox-nnet and DeepSurv, and as the encoder output for AESurv.

For the benchmark methods to compare the performance with BioExPL, we examined two straightforward schemes: (1) feature filtering and (2) feature weighting (Dinh and Ho, 2020). The feature filtering approach reduced the feature dimension by restricting knowledge-guided feature sets only as input to the model. Feature weighting applied the parameter-level L2 regularization to penalize weights associated with non-knowledge features, while leaving knowledge-guided features unregularized. For feature weighting, the regularization strength was varied over 10^0^, 10^−1^, 10^−2^, 10^−3^, 10^−4^. For BioExPL, we set the knowledge alignment hyperparameter *λ* ∈ {10^0^, 10^−1^, 10^−2^, 10^−3^, 10^−4^} as well. For both feature filtering and BioExPL, we optimized the hyperparameters that minimized the validation loss.

#### 3.2.1 BioExPL improves the predictive performance

We conducted 5-fold cross-validation and repeated the evaluation 30 times, reporting the average concordance index (C-index) across all folds and repetitions. BioExPL consistently improved the predictive performance with all the three baseline models across all ten cancer datasets (Fig. 3, Table 2). Overall, the knowledge-guided models (i.e., Cox-nnet, DeepSurv, and AESurv coupled with BioExPL) achieved average C-index values of 0.6684±0.0283, 0.6678±0.0284, and 0.6693±0.0284, respectively, corresponding to improvements of 1.9%, 3.0%, and 1.8% over their baseline performances (0.6557±0.0305, 0.6481±0.0308, and 0.6577±0.0304). BioExPL also outperformed feature filtering and feature weighting schemes. For instance, Cox-nnet with BioExPL achieved the highest C-index of 0.6684*±*0.0283, followed by feature weighting (0.6587*±* 0.0303), the baseline model (0.6557±0.0305), and feature filtering (0.6416±0.0254). BioExPL also showed the highest C-index for DeepSurv and AESurv. The superior performance of BioExPL was statistically validated using the Wilcoxon rank-sum test in the majority of comparisons (*p <* 0.05).

**Table 2.** Predictive performance comparison across ten TCGA cancer datasets.

| Dataset | Cox-nnet |  |  |  | DeepSurv |  |  |  | AESurv |  |  |  |
| --- | --- | --- | --- | --- | --- | --- | --- | --- | --- | --- | --- | --- |
|  | Baseline | Filtering | Weighting | BioExPL | Baseline | Filtering | Weighting | BioExPL | Baseline | Filtering | Weighting | BioExPL |
| LIHC | 0.6319<br>(0.0007) | <u>0.6431</u><br>(0.0018) | 0.6372<br>(0.0005) | **0.6533\*\***<br>(**0.0014**) | 0.6253<br>(0.0015) | <u>0.6480</u><br>(0.0026) | 0.6388<br>(0.0015) | **0.6553\*\***<br>(**0.0016**) | 0.6362<br>(0.0006) | **0.6537\*\***<br>(**0.0012**) | 0.6389<br>(0.0008) | <u>0.6474</u><br>(0.0010) |
| BRCA | 0.6458<br>(0.0013) | 0.6188<br>(0.0028) | <u>0.6479</u><br>(0.0012) | **0.6499**<br>(**0.0011**) | 0.6142<br>(0.0020) | 0.6115<br>(0.0031) | <u>0.6234</u><br>(0.0018) | **0.6390\*\***<br>(**0.0015**) | <u>0.6529</u><br>(0.0010) | 0.6328<br>(0.0012) | 0.6464<br>(0.0012) | **0.6664\*\***<br>(**0.0010**) |
| COAD | 0.5944<br>(0.0014) | 0.5944<br>(0.0044) | <u>0.6034</u><br>(0.0012) | **0.6347\*\***<br>(**0.0027**) | 0.5823<br>(0.0028) | 0.5779<br>(0.0058) | <u>0.5950</u><br>(0.0025) | **0.6292\*\***<br>(**0.0028**) | 0.5961<br>(0.0010) | <u>0.6120</u><br>(0.0018) | 0.5975<br>(0.0007) | **0.6348\*\***<br>(**0.0014**) |
| LUSC | 0.4825<br>(0.0008) | **0.5176**<br>(**0.0020**) | 0.4830<br>(0.0008) | <u>0.5140</u><br>(0.0029) | 0.4858<br>(0.0012) | <u>0.5175</u><br>(0.0033) | 0.4898<br>(0.0012) | **0.5249**<br>(**0.0039**) | 0.4821<br>(0.0006) | <u>0.5026</u><br>(0.0017) | 0.4827<br>(0.0007) | **0.5127\*\***<br>(**0.0028**) |
| SKCM | 0.6262<br>(0.0005) | 0.6314<br>(0.0016) | <u>0.6327</u><br>(0.0005) | **0.6352\***<br>(**0.0016**) | 0.6190<br>(0.0012) | <u>0.6310</u><br>(0.0018) | 0.6275<br>(0.0012) | **0.6322**<br>(**0.0014**) | 0.6279<br>(0.0006) | **0.6371**<br>(**0.0009**) | 0.6300<br>(0.0004) | <u>0.6362</u><br>(0.0007) |
| LUAD | 0.6282<br>(0.0008) | 0.6211<br>(0.0021) | <u>0.6306</u><br>(0.0005) | **0.6346\*\***<br>(**0.0010**) | <u>0.6255</u><br>(0.0010) | 0.6243<br>(0.0017) | 0.6253<br>(0.0009) | **0.6343\*\***<br>(**0.0017**) | 0.6290<br>(0.0006) | **0.6312**<br>(**0.0011**) | 0.6281<br>(0.0006) | <u>0.6301</u><br>(0.0009) |
| LAML | 0.6156<br>(0.0008) | 0.5647<br>(0.0053) | <u>0.6170</u><br>(0.0010) | **0.6187**<br>(**0.0009**) | 0.6022<br>(0.0017) | 0.5792<br>(0.0031) | <u>0.6071</u><br>(0.0018) | **0.6101**<br>(**0.0028**) | <u>0.6209</u><br>(0.0008) | 0.6208<br>(0.0019) | 0.6204<br>(0.0008) | **0.6215**<br>(**0.0009**) |
| KIRC | 0.6943<br>(0.0006) | 0.6642<br>(0.0015) | **0.6970**<br>(**0.0005**) | <u>0.6962</u><br>(0.0009) | 0.6956<br>(0.0009) | 0.6653<br>(0.0018) | <u>0.6982</u><br>(0.0009) | **0.7009\***<br>(**0.0007**) | <u>0.6957</u><br>(0.0007) | 0.6652<br>(0.0008) | 0.6951<br>(0.0006) | **0.6963**<br>(**0.0005**) |
| KIRP | <u>0.7974</u><br>(0.0008) | 0.7542<br>(0.0030) | 0.7957<br>(0.0009) | **0.8018\*\***<br>(**0.0010**) | 0.7963<br>(0.0011) | 0.7323<br>(0.0043) | <u>0.7994</u><br>(0.0016) | **0.8080\*\***<br>(**0.0019**) | <u>0.7944</u><br>(0.0008) | 0.7522<br>(0.0016) | 0.7935<br>(0.0009) | **0.8000\*\***<br>(**0.0010**) |
| GBM/LGG | 0.8403<br>(0.0003) | 0.8061<br>(0.0009) | <u>0.8429</u><br>(0.0004) | **0.8461\*\***<br>(**0.0004**) | 0.8350<br>(0.0004) | 0.8079<br>(0.0008) | <u>0.8392</u><br>(0.0004) | **0.8442\*\***<br>(**0.0005**) | 0.8421<br>(0.0003) | 0.8122<br>(0.0006) | <u>0.8424</u><br>(0.0003) | **0.8479\*\***<br>(**0.0004**) |

Note: reported values indicate the average C-index across repeated evaluations, with standard errors shown in parentheses. For each dataset and model, the best-performing method is highlighted in bold, and the second-best method is underlined. The performance improvements of the best method were further statistically assessed using the Wilcoxon signed-rank test against the second-best method; \* and \*\* denote statistical significance at $p < 0.05$ and $p < 0.01$ , respectively.

**Fig. 3.**
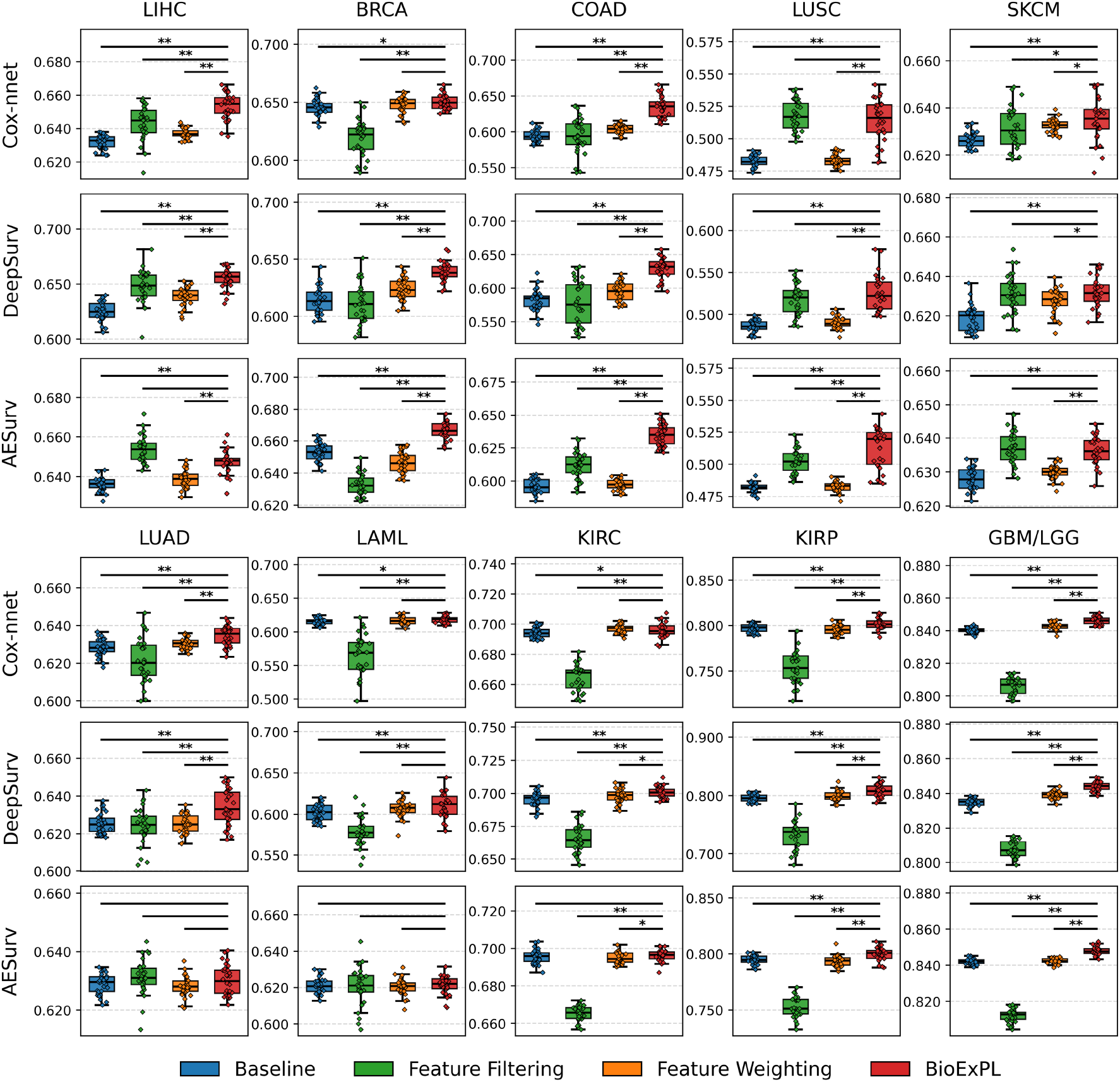
Predictive performance comparison across ten TCGA cancer datasets. Each point represents the average C-index obtained from 5-fold cross-validation. Statistical significance was assessed using the Wilcoxon signed-rank test; ** and * indicate statistically significant improvements with *p <* 0.01 and *p <* 0.05, respectively.

Particularly, BioExPL produced substantial performance gains for LUSC and COAD, with increases of 7.0% and 7.1%, respectively. LUSC has been reported as one of the most challenging cancer types in previous survival analysis studies (Ching et al., 2018; Huang et al., 2020; Meng et al., 2022), often exhibiting C-index values close to 0.5. The observed improvement indicates that BioExPL mitigates this challenge by guiding the model with curated gene associations. COAD is known to be governed by a small number of dominant biological programs that largely influence patient prognosis, as consistently reported by largescale transcriptomic stratification studies (Joanito et al., 2022), pathway-oriented subtype analyses (Komor et al., 2018), and tumor microenvironment–focused investigations (Qi and Zhang, 2022). This molecular characteristic provides a favorable setting for leveraging curated genes, resulting in more pronounced performance improvements over baseline models.

Furthermore, we observed that performance was increased approximately in proportion to the number of guide-associated genes (Fig. 4). The number of curated biomarkers exhibited statistically significant Pearson correlations with performance improvement, with correlation coefficients of 0.64 (*p <* 0.086), 0.98 (*p <* 0.01), and 0.88 (*p <* 0.01) for Cox-nnet, DeepSurv, and AESurv, respectively. These results may imply that the effectiveness of BioExPL is closely related to the available quantity of the prior knowledge set, so baseline models can be further improved by using richer curated biomarkers.

**Fig. 4.**
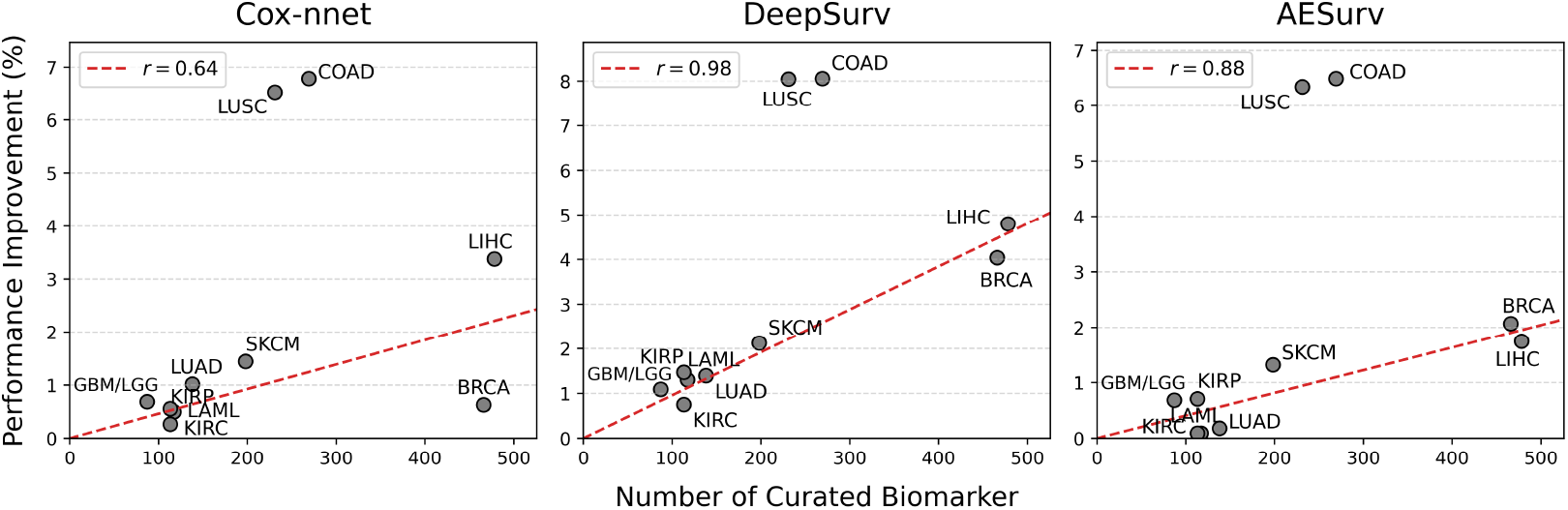
Association between the number of curated biomarkers and the performance improvement achieved by BioExPL across TCGA cancer datasets. Each point represents one cancer cohort, where the x-axis denotes the number of curated biomarkers and the y-axis indicates the relative performance improvement in C-index (%). The red dashed line shows the linear regression fit, and the Pearson correlation coefficient (*r*) is reported. The linear regression and Pearson correlation were estimated excluding the outlier cohorts LUSC and COAD, which exhibited particularly substantial performance improvements.

#### 3.2.2. BioExPL enhances model interpretability

We evaluated whether BioExPL improves the interpretability of the baseline models. Specifically, we examined the top-100 ranked genes by feature importance in the LIHC dataset that contained the largest number of knowledge-guided genes, and compared them with curated gene–disease association sets. Across all baseline models, BioExPL consistently enhanced the model interpretability, identifying more numbers of curated biomarkers among the top-ranked genes (Fig. 5). The knowledge-guided models identified 45, 61, and 62 curated genes among the top-100 genes, respectively, whereas the corresponding baseline models identified only 24, 16, and 22 curated genes each.

**Fig. 5.**
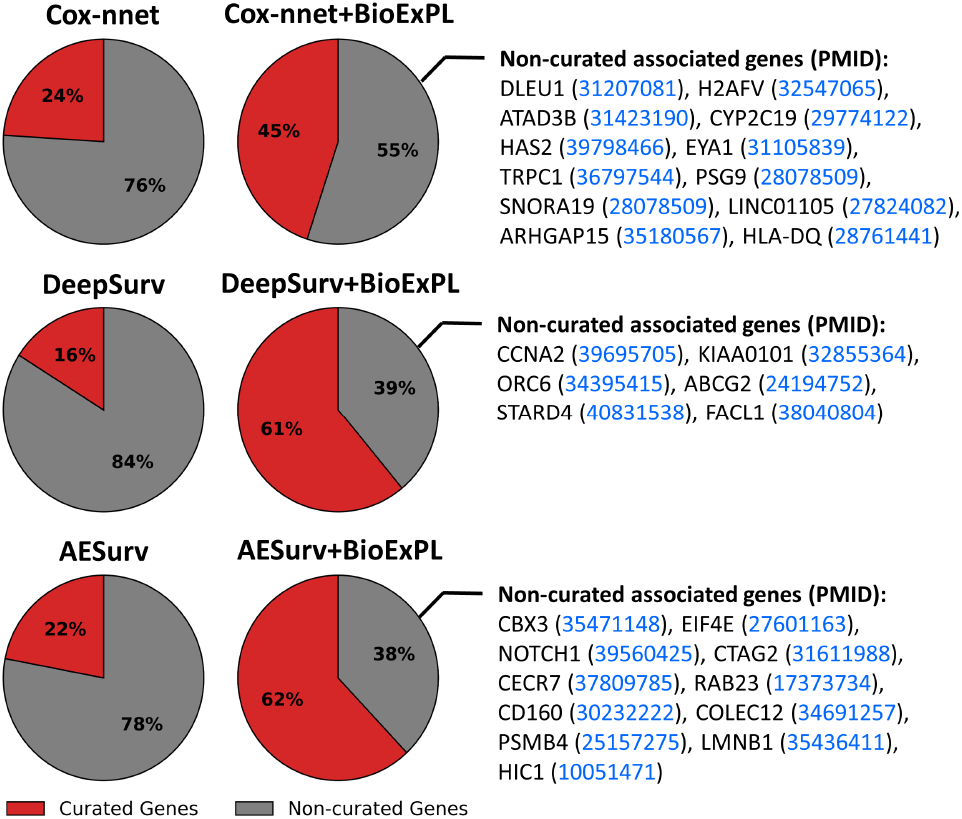
Proportion of curated and non-curated genes among the top 100 features identified by baseline and BioExPL-guided survival models on the LIHC dataset. Top-ranked genes were determined based on gradient-based feature importance scores averaged across repeated evaluations. Non-curated associated genes refer to genes that are not included in the curated biomarker set but were identified by BioExPL; annotated genes are supported by biological literature, with corresponding PubMed identifiers (PMIDs) shown.

We examined whether knowledge-guided models can identify potentially associated genes beyond the curated gene set. We searched the top-ranked genes that were not part of the curated sets from the biological literature and found that several of these genes were reported in independent studies supporting their association with LIHC. Specifically, knowledge-guided Cox-nnet identified 12 additional associated genes among 55 uncurated genes, while knowledge-guided DeepSurv and AESurv identified 7 out of 39 and 11 out of 38 uncurated genes, respectively.

### 3.3 Comparative experiment with random gene sets

We additionally examined whether the performance gains of BioExPL are primarily driven by curated biomarkers, rather than to other potential confounding factors. We hypothesized that substituting the curated gene set with a randomly selected gene set would substantially impair the predictive performance of BioExPL. For each cancer set, we replaced the curated gene set with randomly generated gene set. As a result, BioExPL learning with randomly defined gene sets led to a significant decrease in predictive performance across all cancer datasets (Table 3; *p <* 0.01 in the majority of comparisons). In particular, the average C-index was even lower than that of the corresponding baseline models in the datasets SKCM, LAML, KIRC and KIRP.

**Table 3.** Results of comparative experiments using random gene sets.

| Dataset | Cox-nnet |  | DeepSurv |  | AESurv |  |
| --- | --- | --- | --- | --- | --- | --- |
|  | BioExPL with random genes | BioExPL | BioExPL with random genes | BioExPL | BioExPL with random genes | BioExPL |
| LIHC | 0.6380<br>(0.0011) | **0.6533\*\***<br>(**0.0014**) | 0.6338<br>(0.0016) | **0.6553\*\***<br>(**0.0016**) | 0.6387<br>(0.0008) | **0.6474\*\***<br>(**0.0010**) |
| BRCA | 0.6471<br>(0.0012) | **0.6499\***<br>(**0.0011**) | 0.6248<br>(0.0018) | **0.6390\*\***<br>(**0.0015**) | 0.6593<br>(0.0010) | **0.6664\*\***<br>(**0.0010**) |
| COAD | 0.6208<br>(0.0024) | **0.6347\*\***<br>(**0.0027**) | 0.6000<br>(0.0032) | **0.6292\*\***<br>(**0.0028**) | 0.6150<br>(0.0013) | **0.6348\*\***<br>(**0.0014**) |
| LUSC | 0.5107<br>(0.0038) | **0.5140**<br>(**0.0029**) | 0.5212<br>(0.0027) | **0.5249**<br>(**0.0039**) | 0.5099<br>(0.0024) | **0.5127**<br>(**0.0028**) |
| SKCM | 0.6255<br>(0.0015) | **0.6352\*\***<br>(**0.0016**) | 0.6240<br>(0.0010) | **0.6322\*\***<br>(**0.0014**) | 0.6307<br>(0.0010) | **0.6362\*\***<br>(**0.0007**) |
| LUAD | 0.6302<br>(0.0013) | **0.6346\*\***<br>(**0.0010**) | 0.6335<br>(0.0013) | **0.6343**<br>(**0.0017**) | **0.6330**<br>(**0.0011**) | 0.6301<br>(0.0009) |
| LAML | 0.5989<br>(0.0016) | **0.6181\*\***<br>(**0.0016**) | 0.5892<br>(0.0030) | **0.6101\*\***<br>(**0.0028**) | 0.6185<br>(0.0009) | **0.6215\*\***<br>(**0.0009**) |
| KIRC | 0.6937<br>(0.0007) | **0.6962\***<br>(**0.0009**) | 0.6920<br>(0.0007) | **0.7009\*\***<br>(**0.0007**) | 0.6928<br>(0.0006) | **0.6963\*\***<br>(**0.0005**) |
| KIRP | 0.7968<br>(0.0012) | **0.8018\*\***<br>(**0.0010**) | 0.7963<br>(0.0015) | **0.8080\*\***<br>(**0.0019**) | 0.7931<br>(0.0011) | **0.8000\*\***<br>(**0.0010**) |
| GBM/LGG | 0.8449<br>(0.0003) | **0.8461\***<br>(**0.0004**) | 0.8422<br>(0.0004) | **0.8442\*\***<br>(**0.0005**) | 0.8470<br>(0.0003) | **0.8479\***<br>(**0.0004**) |

Note. reported values indicate the average C-index across repeated evaluations, with standard errors shown in parentheses. For each comparison, the higher-performing method is highlighted in bold. Statistical significance was assessed using the Wilcoxon signed-rank test, where \* and \*\* denote $p < 0.05$ and $p < 0.01$ , respectively.

We also investigated the model performance across varying *λ* in the LIHC dataset, which contained the largest number of curated biomarkers (Fig. 6). We observed that predictive performance consistently decreased across all values of *λ* when random gene sets were used, with more pronounced degradation under stronger knowledge alignment (10^−2^ *< λ*). When BioExPL was applied with curated gene sets, Cox-nnet, DeepSurv, and AESurv achieved performance gains of 3.4%, 4.8%, and 1.7%, respectively, on the LIHC dataset. On the other hand, when using randomly defined gene sets, the corresponding performance gains were only 1.0%, 1.4%, and 0.4%, respectively.

**Fig. 6.**
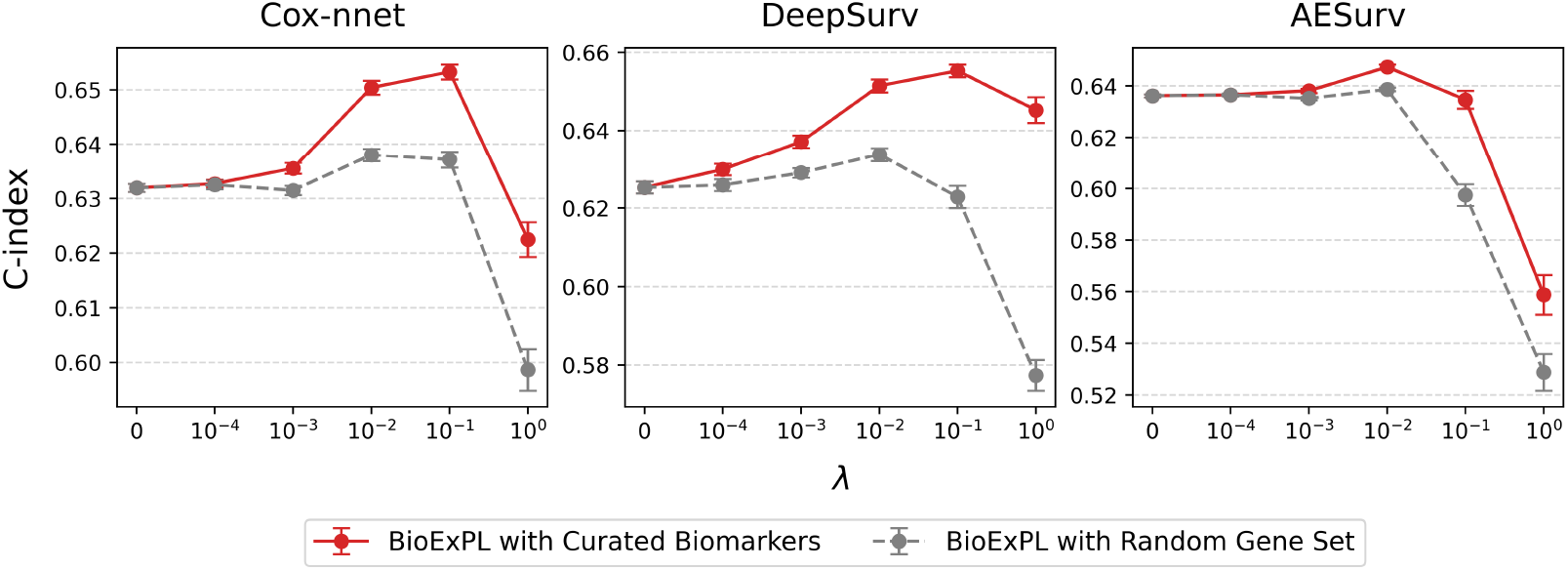
Predictive performance of BioExPL using curated biomarkers and random gene sets on the LIHC dataset. The x-axis represents the knowledge-alignment regularization parameter *λ*. C-index values are shown across varying *λ* for Cox-nnet, DeepSurv, and AESurv.

### 3.4. Computational overhead

We examined additional computational overhead caused by BioExPL, by comparing the average training time per epoch between baseline survival models and their knowledge-guided counterparts. Coupled with BioExPL, Cox-nnet, DeepSurv, and AESurv took 0.0993±0.0002, 0.1001±0.0004, and 0.1131±0.0005 seconds per epoch, respectively (Fig. 7). The corresponding baseline Cox-nnet, DeepSurv, and AESurv models took 0.0925±0.0001, 0.0941±0.0001, and 0.1088±0.0001 seconds per epoch. It resulted in only a modest computational cost of 7.3%, 6.3%, and 4.0% for Cox-nnet, DeepSurv, and AESurv, respectively.

**Fig. 7.**
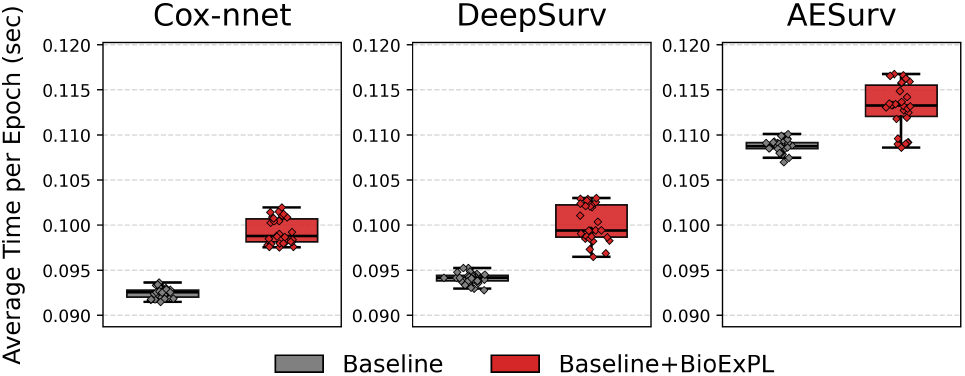
Comparison of computational cost between baseline survival models and their BioExPL-guided counterparts on the LIHC dataset. The average training time per epoch is shown for Cox-nnet, DeepSurv, and AESurv

## 4. Discussion

In this study, we proposed a novel biomarker-driven knowledge-guided learning (BioExPL) by incorporating curated prior knowledge. BioExPL guides neural networks to reflect curated knowledge in their latent representations through a knowledge-alignment loss. Across both simulation studies and large-scale survival analysis, BioExPL consistently improved predictive performance and enhanced model interpretability. In the simulation study, we showed that BioExPL can accurately identify both curated and potentially associated features under controlled settings. In the survival analysis with multiple cancer datasets, BioExPL effectively identified the curated biomarkers and further revealed disease-associated genes supported by external biological literature. Our comparative experiment further supported that the observed gains are driven by meaningful biological priors rather than incidental regularization effects. In terms of efficiency, BioExPL incurred only a modest computational overhead (*<* 7.5%), reflecting an implementation that remains computationally efficient. BioExPL is model-agnostic and domain-independent, enabling its integration into any neural network architectures and application domains.

Despite these overall gains, the effectiveness of BioExPL is still contingent on both the quantity and the quality of the curated knowledge available. In the survival analysis, we observed that BioExPL’s performance improved substantially as more curated biomarkers became available, whereas the gains were relatively modest when only a small set of curated genes was accessible. For instance, BioExPL achieved performance improvements of up to 7% in cancer types with relatively large curated gene sets (e.g., LUSC, COAD, BRCA, and LIHC), while the gains were less than 2% for cancer types with the smallest curated gene sets (e.g., LAML, KIRC, KIRP, and GBM/LGG). In these cancer datasets with smaller curated gene sets, feature filtering led to worse performance than the baseline models, suggesting that the curated sets may not be comprehensive enough to fully represent the underlying disease mechanisms. Moreover, the comparative experiment using random gene set priors demonstrated how the quality of the curated gene set influences the extent of performance improvement.

Curated and experimentally validated knowledge—such as biomarker sets, reference feature sets, and expert-defined catalogs—is among the most common and accessible forms of prior information across scientific disciplines. As a model-agnostic and domain-independent learning strategy, BioExPL can be applied to any neural network-based architectures for high predictive power and robust interpretation with minimum computational burden.

## 5. Acknowledgments

This research was supported by the DOE Office of Science, Office of Biological and Environmental Research (BER) (Grant#: DE-SC0025298) and the National Science Foundation Major Research Instrumentation (NSF MRI) (Grant#: 2117941).

## Footnotes

1 https://ctdbase.org/downloads/#c_gd

